# Glia-specific expression of neuropeptide receptor *Lgr4* regulates development and adult physiology in *Drosophila*

**DOI:** 10.1101/2022.10.07.511249

**Authors:** Seung Gee Lee, Jasdeep Kaur, Hongyu Miao, Woo Jae Kim

## Abstract

Similar to the human brain, *Drosophila* glia may well be divided into several subtypes that each carry out specific functions. Glial GPCRs plays key roles in crosstalk between neurons and glia. *Drosophila Lgr4 (dLgr4*) is a human relaxin receptor homolog involved in angiogenesis, cardiovascular regulation, collagen remodeling, and wound healing. Recent study suggests that *ilp7* might be the ligand for *Lgr4* and regulates escape behavior of *Drosophila* larvae. Here we demonstrate that *Drosophila Lgr4* expression in glial cells, not neurons, is necessary for early development, adult behavior, and lifespan. Reducing the *Lgr4* level in glial cells, but not neurons, disrupts *Drosophila* development, although knocking down other LGR family members in glia has no impact. Adult-specific knockdown of *Lgr4* in glia but not neurons reduce locomotion, male reproductive success, and animal longevity. The investigation of how glial expression of *Lgr4* contributes to this behavioral alteration will increase our understanding of how insulin signaling via glia selectively modulates neuronal activity and behavior.

## INTRODUCTION

Neurons and glia are the two main cell types found in nervous systems. Glia has long been recognized as playing a significant role in regulating the survival of neurons in both vertebrates and invertebrates [1]. Glial cells are necessary for the normal development and functioning of the nervous system. Glia engulfs extraneous cells and projections throughout development to control the number of neuronal cells and guarantee that mature brain circuits are properly shaped. Glial cells in the adult brain provide crucial immunological support and metabolic support in response to short-term and long-term challenges. Age-related cognitive decline and the risk of neurodegenerative diseases are thought to be influenced by dysfunctional glial immune activation [2–5].

The structure and function of glia in *Drosophila* are very similar to those in mammals [6]. Similar to the human brain, *Drosophila* glia may well be divided into several subtypes that each carry out specific functions, such as CNS pruning, controlling synaptic signals, forming the blood-brain barrier, and providing neuroprotection. Recent studies in *Drosophila* suggest that glia supports the systemic metabolism for brain health [7,8]. For instance, glial glycolysis, which is mediated by glycolytic enzymes mostly produced by blood-brain barrier (BBB) glia, is crucial for neuronal survival and is involved in mediating sugar import into the *Drosophila* brain [9].

G-protein coupled receptors (GPCRs) are the largest superfamily of membrane proteins. It is believed that almost 50% of medicines target GPCRs [10]. Glial GPCRs plays key roles in crosstalk between neurons and glia [11], modulation of synaptic transmission at the neuromuscular junction [12], adult neurogenesis [13], astrocyte-neuron communication [14], and BBB formation [15,16]. In *Drosophila*, the expression of GPCR *Moody* in subperineurial glia is critical in order to maintain the integrity of BBB and modulate cocaine, nicotine and ethanol sensitivity [15,17–19].

Leucine-rich repeat containing G protein-coupled receptors (LGRs) make up a group of transmembrane proteins that are distinguished by having a large N-terminal extracellular domain. *Drosophila Lgr4 (dLgr4*) shares 40% amino acid similarity with human relaxin receptors [20]. Mammalian *relaxins* and their receptor signaling are involved in a variety of activities, including angiogenesis, cardiovascular control, collagen remodeling, and wound healing [21]. *Lgr4* is strongly expressed in the adult male brain, the thoracic-abdominal ganglion, the crop, and particularly in the male midgut [22], in contrast to *Lgr3*, which is highly expressed in larval CNS and regulates the timing of developmental transitions through its ligand *ilp8* binding [23,24]. Recent study suggests that *ilp7* might be the ligand for *Lgr4* and *ilp7-Lgr4* neuropeptidergic circuit gates selective escape behavior of *Drosophila* larvae [25].

In this study, we demonstrated that *Drosophila Lgr4* expression in glial cells, not neurons, is necessary for early development, adult locomotion, and lifespan.

## Results

### Glial knockdown of *Lgr4* disrupt the development of *Drosophila*

To investigate the role of *Lgr4* in development, we used pan-glial *repo-GAL4* and pan-neuronal *elav^c155^* drivers to knock-down *Lgr4*. Control experiments for the lethality testing revealed that in both males and females, the ratio of eclosed animals with TM3 balancer chromosomes and wild-type chromosomes is similar in all temperatures (Fig. 1A-B and Fig. S1A-B). Flies that expressed *Lgr4-RNAi* with *repo-GAL4* showed severe mortality when reared at 25°C and 29°C (Fig. 1C-D), but those that expressed with *elav^c155^* developed normally at all temperatures (Fig. 1E-F). Males showed a more severe lethality phenotype than females when *Lgr4* is knocked down in glial cells (compare the 29°C bars between Fig. 1C and 1D). We verified three distinct *Lgr4-RNAi* strains and concluded that *repo-GAL4* is effective with all three RNAi strains (Fig. S1C-D). All these data suggest that glial expression of *Lgr4* is essential for normal development in *Drosophila*.

**Fig. 1.**
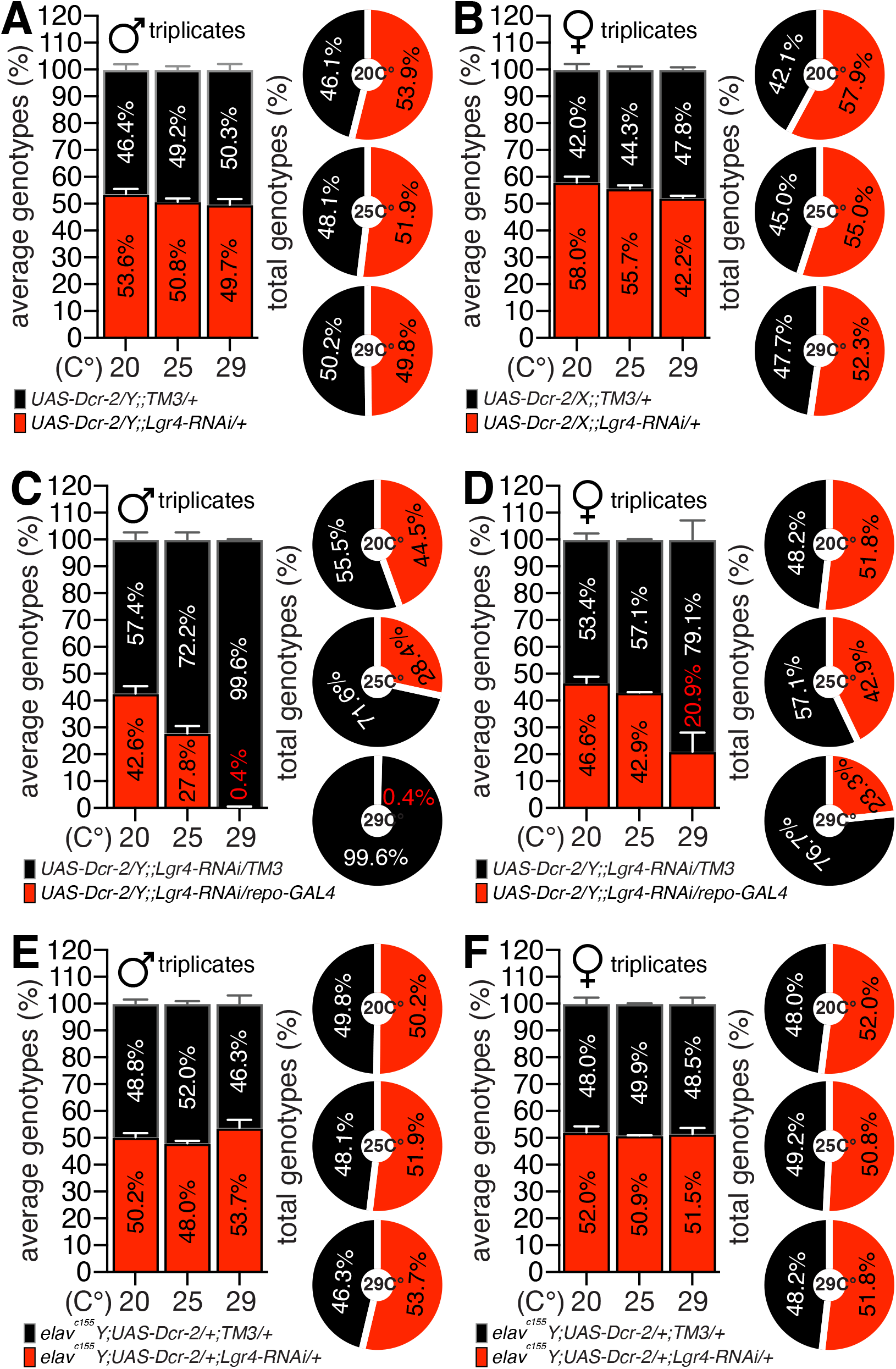
The impact of *Lgr4* knockdown on neuronal and glial cell populations, as determined by the percentage of lethality from egg to adult. (A-B) Experiments serving as controls for the lethality test used in this study. The proportion of (A) male or (B) female flies with the *TM3* balancer chromosome (black bar) or *RNAi* (red bar) is shown in the left bar graph. All results were collected from the triplicate vials. The right donut graph represents the accumulated data from the triplicate tests shown on the left bar graph. (C-D) The average (left bar) and accumulated (right donut) proportion of (C) male or (B) female flies with *TM3* balancer chromosome (black) or *RNAi* (red) conjugated with *repo-GAL4* driver. (E-F) The average (left bar) and accumulated (right donut) proportion of (E) male or (F) female flies with *TM3* balancer chromosome (black) or *RNAi* (red) conjugated with *elav^c155^* driver. Sex of each animal is labeled above the graph. Genotypes of each animal is labeled below the graph. The numbers inside the bar or donut graphs represent the proportion of an eclosed animal. See **EXPERIMENTAL PROCEDURES** for detailed experimental methods and statistical analysis used in this study.

*Drosophila* genome contains four LGRs family, *Lgr1, Lgr2, Lgr3*, and *Lgr4*. Fly *Lgr1* belongs to type A LGRs, which contain mammalian glycoprotein hormone receptors (FSHR, LH/CGR, and TSHR). Fly *Lgr2* belongs to type B LGRs, which contain mammalian LGR4, 5, and 6. Fly *Lgr3* and *Lgr4* belong to type C LGRs and contain mammalian LGR7 and LGR8, also called RXFP1 and RXFP2, respectively [20]. In order to compare the role of other fly LGRs family with *Lgr4* in glial cells, we expressed *Lgr1-, Lgr2, Lgr3-RNAi* using *repo-GAL4*. Male and female flies that expressed *Lgr1-RNAi, Lgr2-RNAi*, or *Lgr3-RNAi* with *repo-GAL4* didn’t exhibit any abnormalities in the lethality test (Figs. 2A-F).

**Fig. 2.**
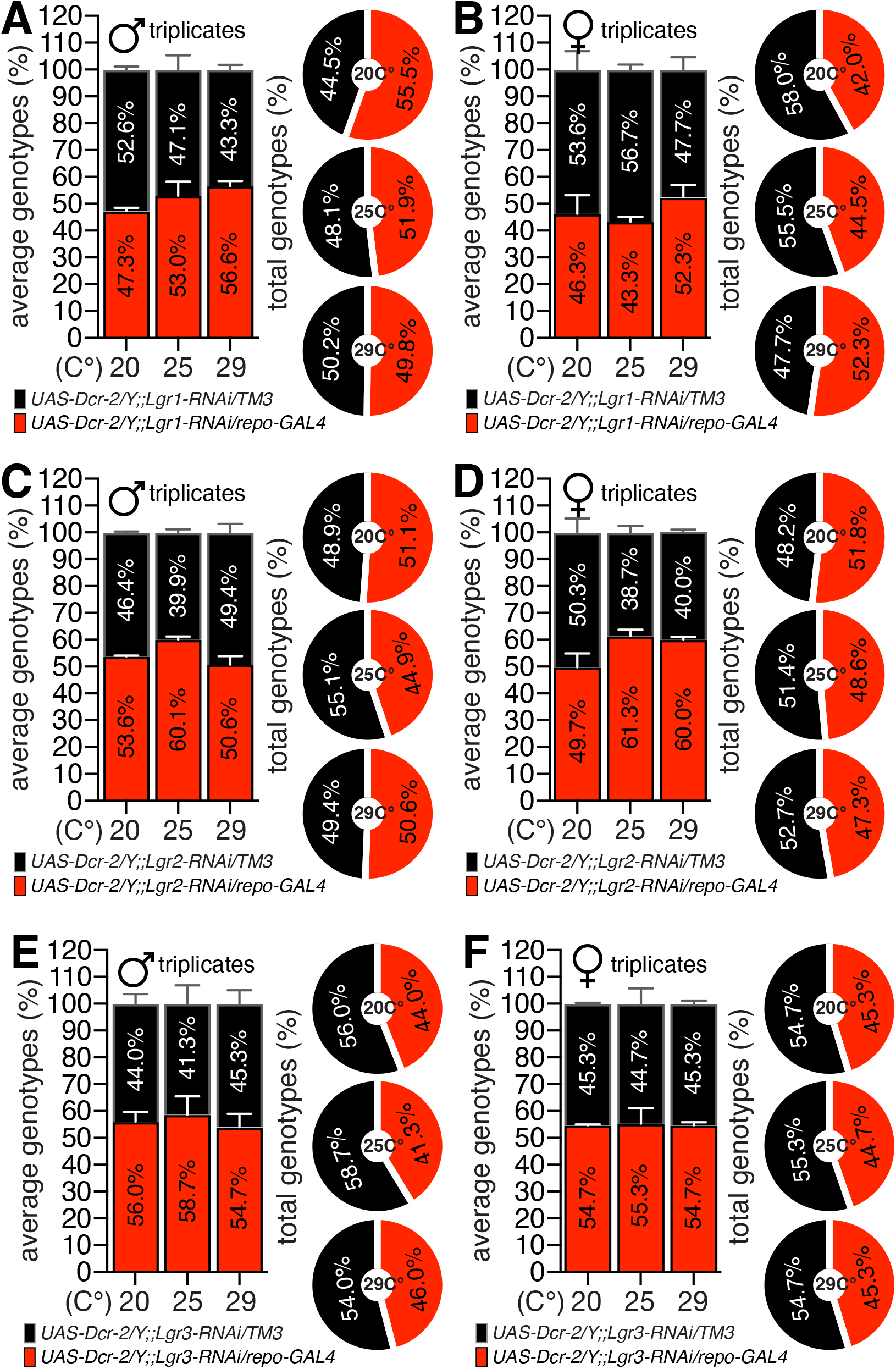
The impact of LGRs knockdown on glial cell populations, as determined by the percentage of lethality from egg to adult. (A-F) The average (left bar) and accumulated (right donut) proportion of (A, C and E) male or (B, D and F) female flies with *TM3* balancer chromosome (black) or *repo-GAL4* (red) conjugated with (A-B) *Lgr1-RNAi* (C-D) *Lgr2-RNAi* (E-F) *Lgr3-RNAi*. Sex of each animal is labeled above the graph. Genotypes of each animal is labeled below the graph. The numbers inside the bar or donut graphs represent the proportion of an eclosed animal.

Next, we confirm previously described glia-*GLA4s* drivers (Table. 1) using *Lgr4-RNAi* to determine which subtype glia is associated with these developmental consequences. Only animals expressing *Lgr4-RNAi* in conjunction with *Eaat1-* and *Mz97-GAL4* drivers exhibited a lethal phenotype among those commonly used glial-*GAL4* drivers (Fig. 3A-B). Similar to the results of *repo-GAL4*, males showed a more severe lethality phenotype than females (*Eaat1^8849^-GAL4:* 4.1% vs. 26.7%, *Eaat1^P88^-GAL4:* 18.2% vs. 22.5%, and *Mz97-GAL4:* 13.1% vs. 22.5%). Next, we used recently developed subtype glial-*GAL4* drivers to determine which subtypes of glia is responsible for this phenotype. Both males and females expressing *Lgr4-RNAi* conjugated with *ALG-, CG-, PNG-*, and *SPG-GAL4* drivers had a lethal phenotype, with males exhibiting a more severe phenotype than females (Fig. 3C-D), as shown by *repo-GAL4* (Fig. 1C-D) and other glial-*GAL4* drivers (Fig. 3A-B). All of these findings show that the expression of Lgr4 in certain subtypes of glia is crucial for proper development.

**Fig. 3.**
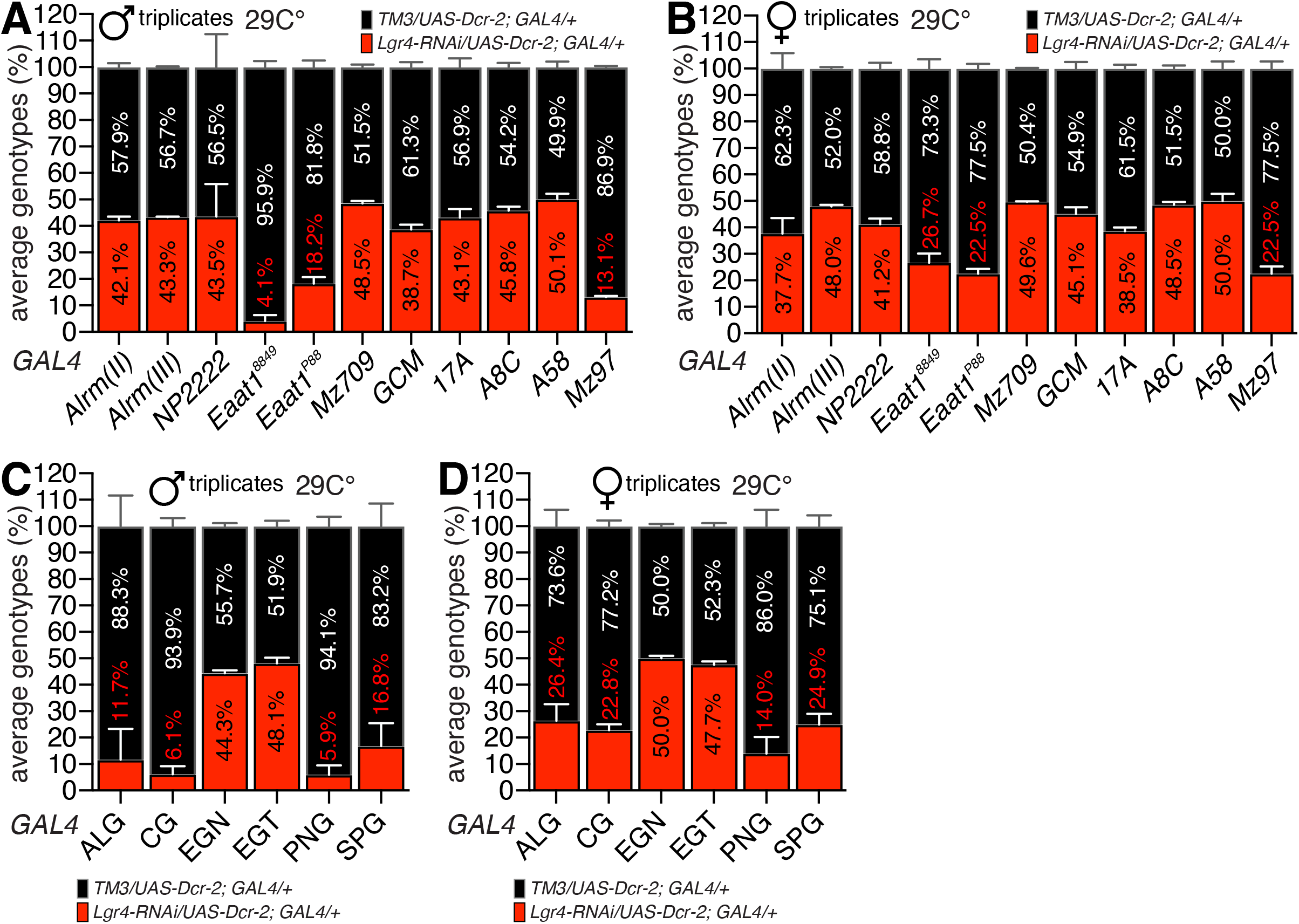
Testing the functionality of glial *GAL4* drivers with *Lgr4-RNAi*. (A-B) The average (left bar) proportion of (A) male or (B) female flies with *TM3* balancer chromosome (black) or *repo-GAL4* (red) conjugated with (A-B) *Lgr1-RNAi* (C-D) *Lgr2-RNAi* (E-F) *Lgr3-RNAi*. Sex of each animal is labeled above the graph. Genotypes of each animal is labeled below the graph. The numbers inside the bar or donut graphs represent the proportion of an eclosed animal.

**Table 1.** Description of glial *GAL4* drivers used in this study.

| Name | genotype | Ch'me | Expression | References | Stock # |
| --- | --- | --- | --- | --- | --- |
| *Alrm* | *P{alrm-GAL4.D}2* | II | astrocyte-like glial cell | (Doherty 2009) | BDSC 67032 |
|  | *P{alrm-GAL4.D}3* | III | adult optic chiasma giant glial cell |  | BDSC 67031 |
| *NP2222* | *P{GawB}Akap200<sup>NP2222</sup>* | II | cell body glial cell<br>cortex glia | (Lee 2019) | Kyoto 112830 |
| *Eaat1<sup>8849\*</sup>* | *P{Eaat1-GAL4.R}2* | II | astrocytes and other glia | (Rival 2004) | BDSC 8849 |
| *Eaat1<sup>P88\*</sup>* | *P{Eaat1-GAL4.R}P88* | II | astrocytes and other glia | (Rival 2004) | N/A |
| *Mz709* | *P{GawB}Mz709* | N/A | ensheathing glia | (Doherty 2009) | N/A |
| *GCM* | *P{gcm-GAL4.U}* | N/A | neuroglioblast/glioblast-specific | (Hosoya 1995) | N/A |
| *17A* | *P{GawB}17A* | II | brain subperineurial glial cell | (King 1988) | BDSC 8474 |
| *A8C* | *P{GawB}A8C* | III | adult medulla neuropil associated glial cell<br>epithelial glial cell | (Chotard 2005) | BDSC 35543 |
| *A58* | *P{GAL4} A58* | N/A | embryonic larval epidermis | (Galko 2004) | N/A |
| *Mz97\** | *P{GawB}Ntan1<sup>Mz97</sup>* | II | surface glial cell<br>wrapping glial cell<br>lamina anlage glial cell | (Cameron 2015) | BDSC 9488 |
| *ALG\** | *R86E01-GAL4* (JFRC) | III | astrocyte-like glia | (Kremer 2017) | BDSC 45914 |
| *CG\** | *R77A03-GAL4* (JFRC) | III | cortex glia | (Kremer 2017) | BDSC 39944 |
| *EGN* | *R56F03-GAL4* (JFRC) | III | neuropile ensheathing glia | (Kremer 2017) | BDSC 39157 |
| *EGT* | *R75H03-GAL4* (JFRC) | III | tract ensheathing glia | (Kremer 2017) | BDSC 39908 |
| *PNG\** | *R85G01-GAL4* (JFRC) | III | perineurial glia | (Kremer 2017) | BDSC 40436 |
| *SPG\** | *R54C07-GAL4* (JFRC) | III | subperineurial glia | (Kremer 2017) | BDSC 50472 |
| *repo\** | *P{GAL4}repo* | III | glial cell | (Sepp 2001) | BDSC 7415 |

\* Reflects the *GAL4s* affect the lethality of development when combined with *Lgr4-RNAi*

### Glial knockdown of *Lgr4* in adults *Drosophila* shows severe locomotion and lifespan defects

In order to investigate the role of *Lgr4* in adult stage, we expressed *Lgr4-RNAi* at 20°C throughout development to diminish the effects of RNAi. After adult eclosion, we raise the temperature to 29°C to enhance the effects of RNAi throughout the adult stage [26]. Males and females expressing *Lgr4-RNAi* in association with *repo-GAL4* displayed abnormal locomotion, but those expressing *elav^C155^* exhibited normal locomotion (Fig. 4A-B). Males expressing Lgr4-RNAi in glia failed to mate, whereas those in neurons mated successfully (Fig. 4C). Lack of *Lgr1, Lgr2*, or *Lgr3* in glia was not associated with mating success (Fig. 4D). Only males lacking glial *Lgr4*, but not neuronal *Lgr4*, had a substantially shorter lifetime at 25°C and 29°C (Fig. 4E-G). All of these findings indicate that Lgr4 glial activity is required for appropriate adult locomotion and longevity.

**Fig. 4.**
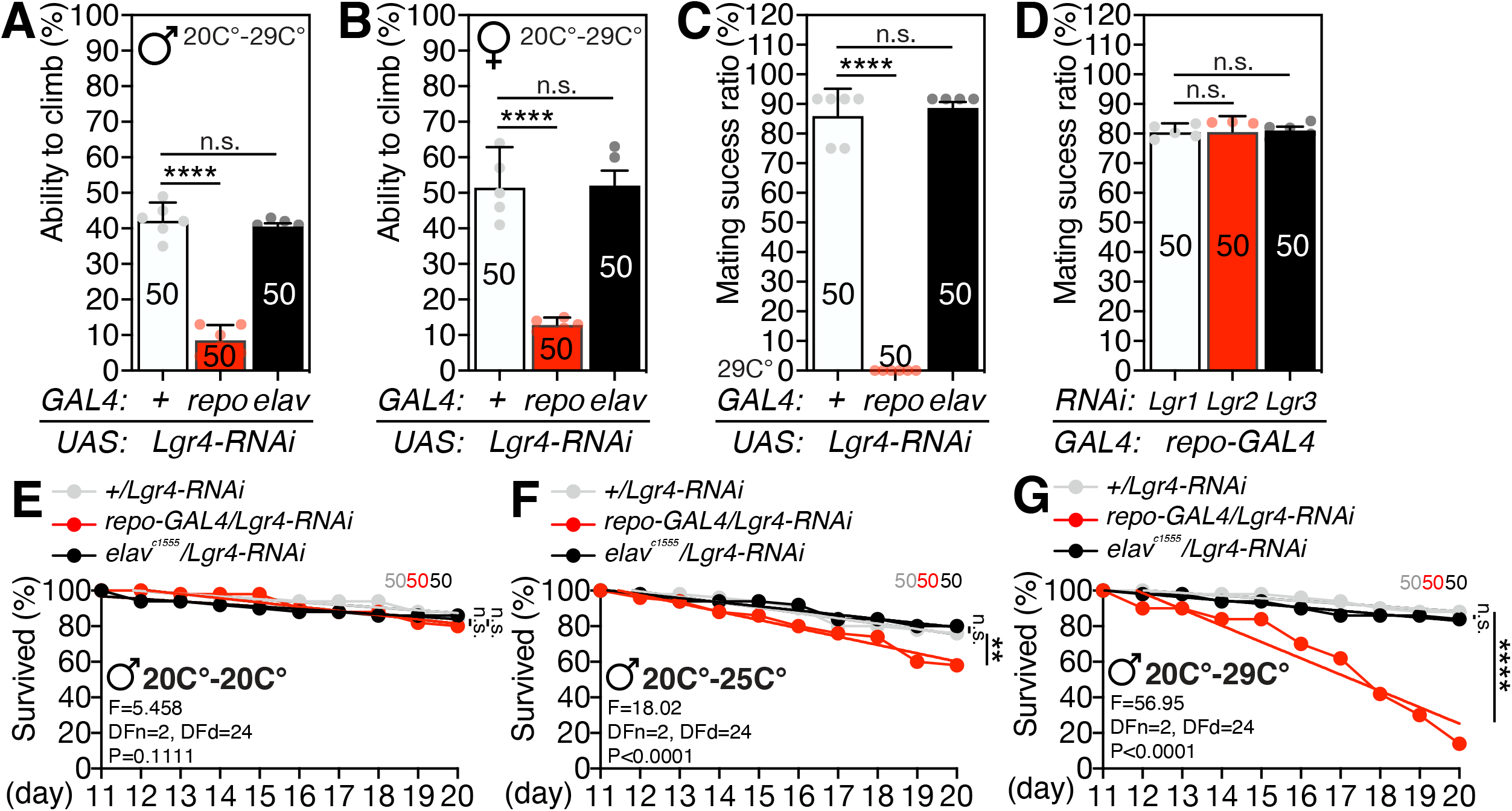
The impact of Lgr4 knockdown in adult behavior and physiology. (A-B) Bar graphs with dot plot represent the percent of (A) male or (B) female flies crossed midline. All experiments were performed five times after sufficient recovering period. (C-D) Bar graphs with dot plot represent the percent of male flies successfully mated within 1 hour with wild-type virgin female. (E-G) Each colored dot represents the percentage of survived flies at that day. Colored slope represents the line by the linear regression analysis. The difference between lines were inferred by linear regression analysis. Genotypes of all above data are labeled below the graph. Rearing temperature, age, and sex of animals are labeled within the graph. Asterisks represent significant differences revealed by unpaired Student’s *t* test (\**p<0.05*, \*\**p<0.01*, \*\*\**p<0.001*, \*\*\*\**p<0.0001*). n.s. represent non-significant differences revealed by unpaired Student’s *t* test. The same notations of climbing assay for statistical analysis are used in other figures. See **EXPERIMENTAL PROCEDURES** and previous report [31,32] for detailed quantification methods of climbing assay, mating assay and lifespan assay.

### *Lgr4* expresses highly in adult glial cells

To validate the glial expression of *Lgr4*, we made use of a scRNA sequencing dataset of fruit fly that is available on the SCope website [27]. Using the annotation function of SCope, we confirmed that adult glial cell highly expresses *Lgr4* both in male and female (Fig. 5A-B). Notably, the newly proposed *Lgr4* ligand, *ilp7*, is not coexpressed with *Lgr4* nor expressed in adult glial cells (Figure 5C-E). In contrast to *Lgr4*, adult glial cells do not express *Lgr3* (Fig. 5G-H). *Lgr4*-expressing cells partly overlap with annotated ‘cell body glia’ and ‘adult glial cell’ when just ‘glial cell’ is considered (Fig. 5I-T). Most *Lgr4*-expressing glial cells were enriched in the “body,” not the “head,” both in males and females (Fig. 5U-X). Male “testis” and “body wall” of both sexes had more *Lgr4*-expressing cells than other identified tissues (Fig. S4). *Lgr4*-expressing cells were enriched in ‘epithelial cell,’ ‘glial cell,’ and ‘muscle’ based on functional annotation (Fig. S5 and Fig. S6A-D).

**Fig. 5.**
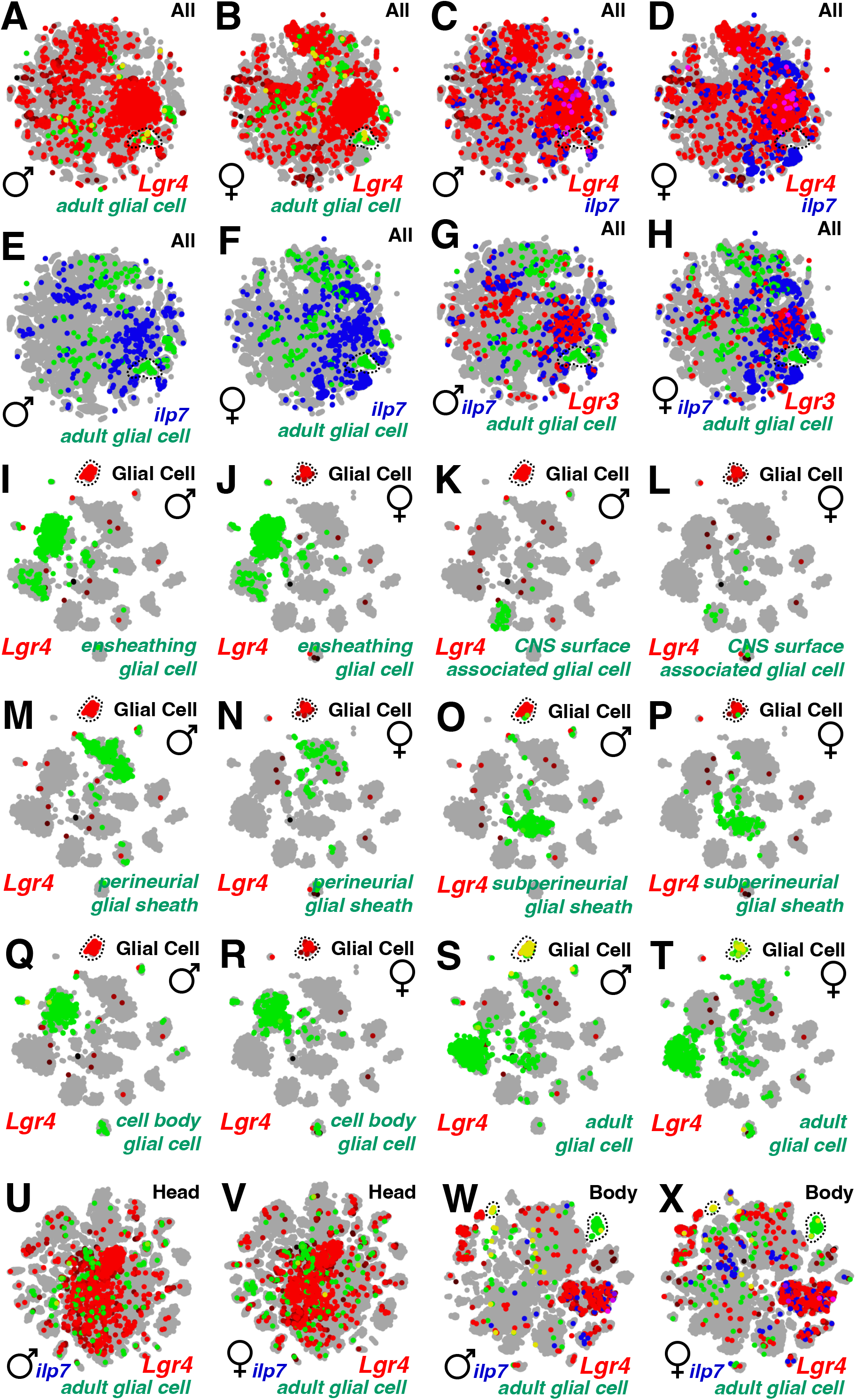
snRNAseq of adult fly [27] confirms the expression of *Lgr4* in glial cells among the various tissues annotated by fly SCope (https://flycellatlas.org/scope). (A-H) SCope data for ‘All’ tissue. (I-T) SCope data for annotated ‘Glial Cell’ populations. (U-V) SCope data for ‘Head’. (W-X) SCope data for ‘Body’. Annotations and gene names of all above data are color-coded using red, green, and blue words. When cells overlap, the color of the dots is either yellow, cyan, or magenta. Dotted lines indicate glial cell populations that expresses *Lgr4*.

To validate the expression patterns of *Lgr4* throughout development and adulthood, we analyzed the expression of existing *Lgr4-GAL4* strains (Fig. 6A). When coupled with *UAS-CD4tdGFP*, Mi{ET1}-based two enhancer trap *GAL4* strains, *Lgr4^MB01485^-GAL4* and *Lgr4^MB03440^-GAL4* expression demonstrated glia patterns in both male and female CNS (Fig. 6B, C, E, and F). The intensity of GFP fluorescence is greater in the VNC than in the brain (Fig. 6D and G). When paired with membrane and nuclear markers, *UAS-mCD8GFP, UAS-RedStinger*, putative brain enhancer *GAL4* strains [28] *R24A08-GAL4* and *R25C08-GAL4* label a huge quantity of glial cells in the larval CNS (Fig. S6E-F and Fig. 7A). When *repo-GAL80* is coupled to *R25C08-GAL4*, membrane and nucleus patterns of glia (Figure 7B-C) disappeared (Fig. 7F-H). *RedStinger* fluorescence quantification revealed that about 85 percent of all cells tagged by *R25C08-GAL4* were eliminated by *repo-GAL80* conjugation (Fig. 7I). *Lgr4MB03440-GAL4* also tagged adult brain surface glia in both sexes (Fig. S6G) as shown in FlyLight surface glia *GAL4s* (Fig. S6I-J), and *repo-GAL80* conjugation could reduce this expression patterns (Fig. S6H). GFP expression was more prevalent in male brains than in female brains (Fig. S6K). *Repo-GAL80* conjugation with *Lgr4^MB03440^-GAL4* lowers GFP expression in the male brain by around 90% (Fig. S6L). These findings indicate that Lgr4 is extensively expressed in glia throughout both the developmental and adult stages.

**Fig. 6.**
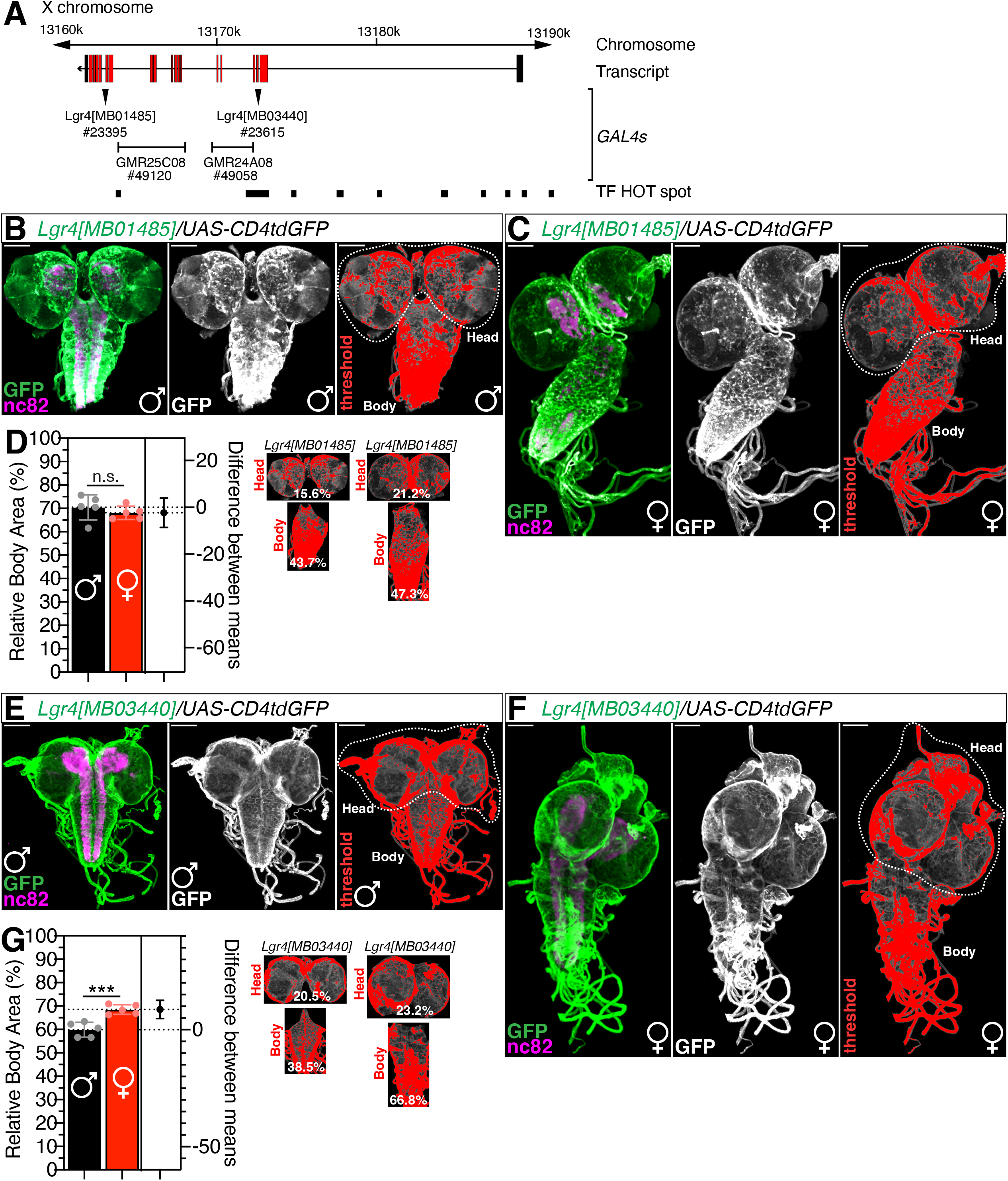
*Lgr4-GAL4s* label glia in larval CNS. (A) Enhancer elements of Lgr4 gene. The genotypes and stock numbers for all four GAL4 drivers are described. TF HOT spot represents the conserved transcription factor binding motifs deduced from ChIP data of whole embryo by REDfly project [33]. Diagram is from *Lgr4* gene region of FlyBase GBrowse. (B-C) Third instar (B) male or (C) female larval CNS expressing *Lgr4[MB014850]* with *UAS-CD4tdGFP* were immunostained with anti-GFP (green) and anti-nc82 (magenta) antibodies. Gray panel represents GFP signals for quantification to set threshold shown in the right panels. Scale bars represent 100 μm. (D) Percent of body area quantified from (B) and (C). GFP signals represents GAL4 expressing ‘Head’ or ‘Body’ region were calculated by ImageJ threshold and measure function then quantified. (E-G) The same experiments were performed as described in (B-D) except the *Lgr4[MB03440]* was used as *GAL4* driver. See **EXPERIMENTAL PROCEDURES** and previous report [31,32] for detailed quantification methods.

## DISCUSSION

Our study shows that *Lgr4* expression in a subset of glial cells is essential for normal development and adult physiology. Reducing the *Lgr4* level in glial cells, but not neurons, disrupts *Drosophila* development (Fig. 1 and S1), although knocking down other LGR family members in glia has no impact (Fig. 2). Using previously reported glial-*GAL4* drivers and newly discovered subtype glia-specific *GAL4* drivers, we determined that ALG, CG, PNG, and SPG are the subtype glial populations responsible for *Lgr4*-mediated normal development (Fig. 3). Adult-specific knockdown of *Lgr4* in glia but not neurons reduce locomotion, male reproductive success, and animal longevity (Fig. 4 and S4). Using newly published *Drosophila* scRNA seq data and accessible *Lgr4-GAL4* drivers, we showed that *Lgr4* is substantially concentrated in adult body glia but not head glia (Fig. 5, S4, S5, 6, S6 and 7).

**Fig. 7.**
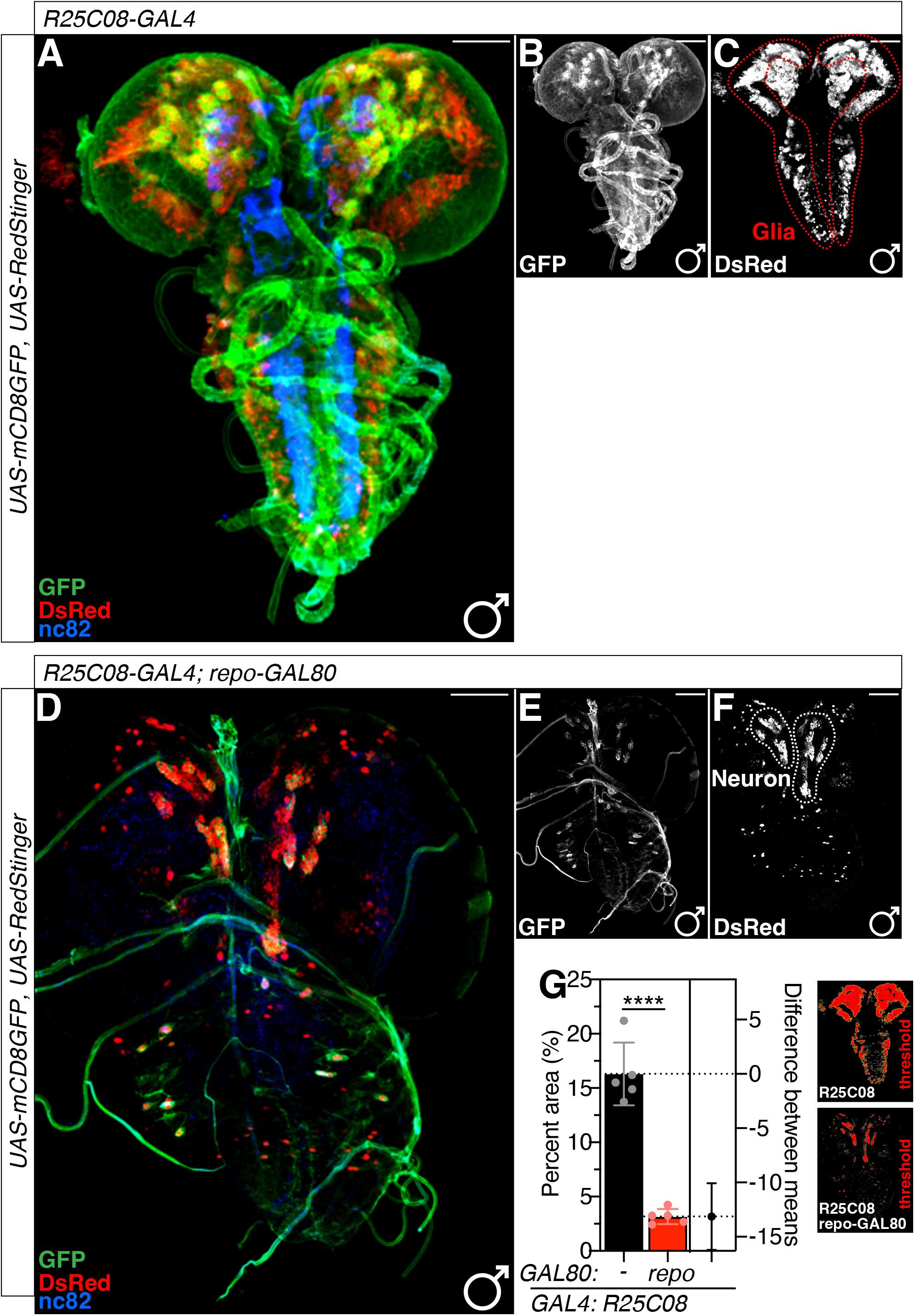
The most of *Lgr4-GAL4-labeled* cells are glia. (A-F) Third instar larval CNS expressing *UAS-mCD8GFP, UAS-RedStinger* with (A-C) *R25C08-GAL4* or (D-F) *R25C08-GAL4, repo-GAL80* were immunostained with anti-GFP (green), anti-DsRed (red) and anti-nc82 (blue) antibodies. Gray panel represents GFP and RedStinger signals for quantification to set threshold shown in (G). Scale bars represent 100 μm. Red dotted lines represent nuclei of glia and white dotted lines nuclei of neurons. (G) Percent area of RedStinger signals quantified from (C) and (F). RedStigner signals represents the number of *GAL4* expressing cells were calculated by ImageJ threshold and measure function then quantified. Right panel represents RedStinger signals of set threshold by ImageJ for quantification. See **EXPERIMENTAL PROCEDURES** for detailed quantification methods.

One of the intriguing features of *Lgr4* function is that males are more sensitive to knockdown of *Lgr4* in glial cells (Fig. 1C and D, Fig. S1C and D, Fig. 3A and B, Fig. 3C and D). It has been shown that the level of *Lgr4* expression in males is much higher than that in females [22]. *Lgr4* transcript is highly expressed in the third instar larvae’s midgut, Malpighian tubules, hindgut, central nervous system, salivary glands, and fat body. The adult stages of the midgut, crop, brain, and thoracic-abdominal ganglia express *Lgr4* transcript at high levels [22]. Mamalian homologues of LGRs are the relaxin-family peptide receptors RXFP1 and RXFP2, which, among other roles, are required for proper reproduction in both sexes [21]. Since Lgr3 in *Drosophila* neurons was reported to be sex-specifically regulated [29] and controls developmental timing with its ligand ilp8 [23,24], it will be interesting to determine whether ilp7-Lgr4 signaling through neuron-glia in *Drosophila* is also sex-specifically regulated.

Relaxin subfamily in vertebrates play diverse roles in tissue homeostasis and remodeling, behaviour and reproduction [21]. Recent reports indicate that ilp7-Lgr4 signaling works as circuit gates for escape behavior [30]. The investigation of how glial expression of *Lgr4* contributes to this behavioral alteration will increase our understanding of how insulin signaling via glia selectively modulates neuronal activity and behavior.

## ACKNOWLEDGEMENTS

We are very appreciative to the colleagues who supplied us with several fly strains; Dr. Marc Freeman (Vollum Institute, OHSU, USA), Drs. Qingzhong Ren and Tzumin Lee (HHMI Janelia Research Campus, USA), Dr. Chun Han (Cornell University, USA) *A58-GAL4*. Stocks obtained from the Bloomington Drosophila Stock Center (NIH P40OD018537) were used in this study. Transgenic fly stocks and/or plasmids were obtained from the Vienna Drosophila Resource Center (VDRC, www.vdrc.at). The fly stock was obtained from KYOTO *Drosophila* Stock Center in Kyoto Institute of Technology. This work was supported by University of Ottawa Startup grant to WJK, University of Ottawa Brain and Mind Research Institute/Center for Neural Dynamics Open call project grant to WJK, University of Ottawa Interdisciplinary Research Group Funding Opportunity (IRGFO stream 1 and 2) Grant to WJK, Natural Sciences and Engineering Research Council of Canada (NSERC) Discovery grant 211406 to WJK, Mitacs Globalink Research Internship Program grant to WJK, and Startup funds from HIT Center for Life Science to WJK. This work was also supported by a Brain Pool Program by National Research Foundation in Korea to WJK, Burroughs Wellcome Fund Collaborative Research Travel Grants 1017486 to WJK, NVIDIA Academic Hardware Grant Program to WJK.

---

**Fig. S1.**
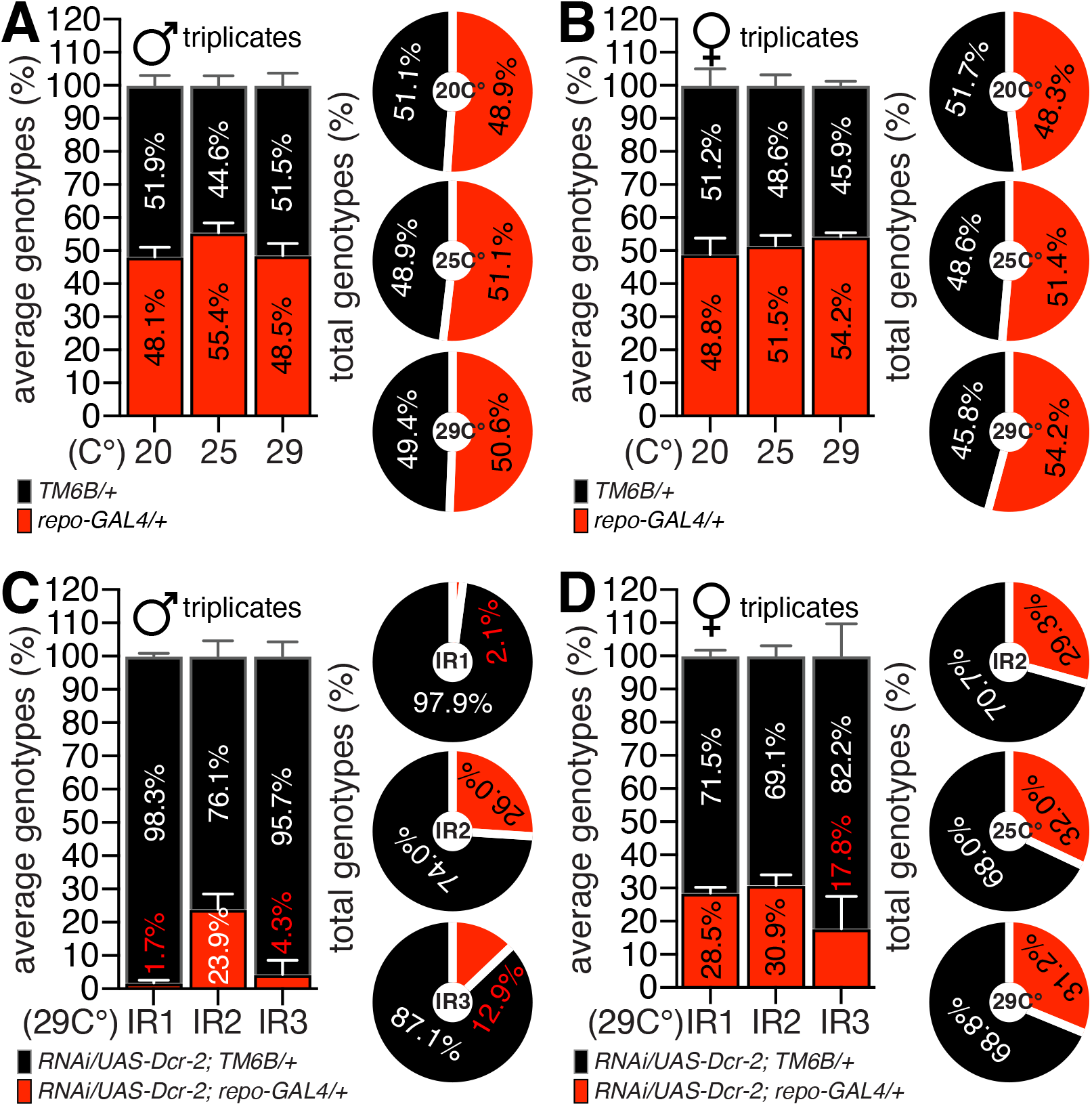
Control experiments for *Lgr4-RNAi* knock down experiments. (A-B) Experiments serving as controls for the lethality test used in this study. The proportion of (A) male or (B) female flies with the *TM6B* balancer chromosome (black bar) or *repo-GAL4* (red bar) is shown in the left bar graph. All results were collected from the triplicate vials. The right donut graph represents the accumulated data from the triplicate tests shown on the left bar graph. (C-D) The average (left bar) and accumulated (right donut) proportion of (C) male or (B) female flies with *TM3* balancer chromosome (black) or *RNAi* (red) conjugated with *repo-GAL4* driver. Genotypes of each animal is labeled below the graph. The numbers inside the bar or donut graphs represent the proportion of an eclosed animal.

**Fig. S3.**
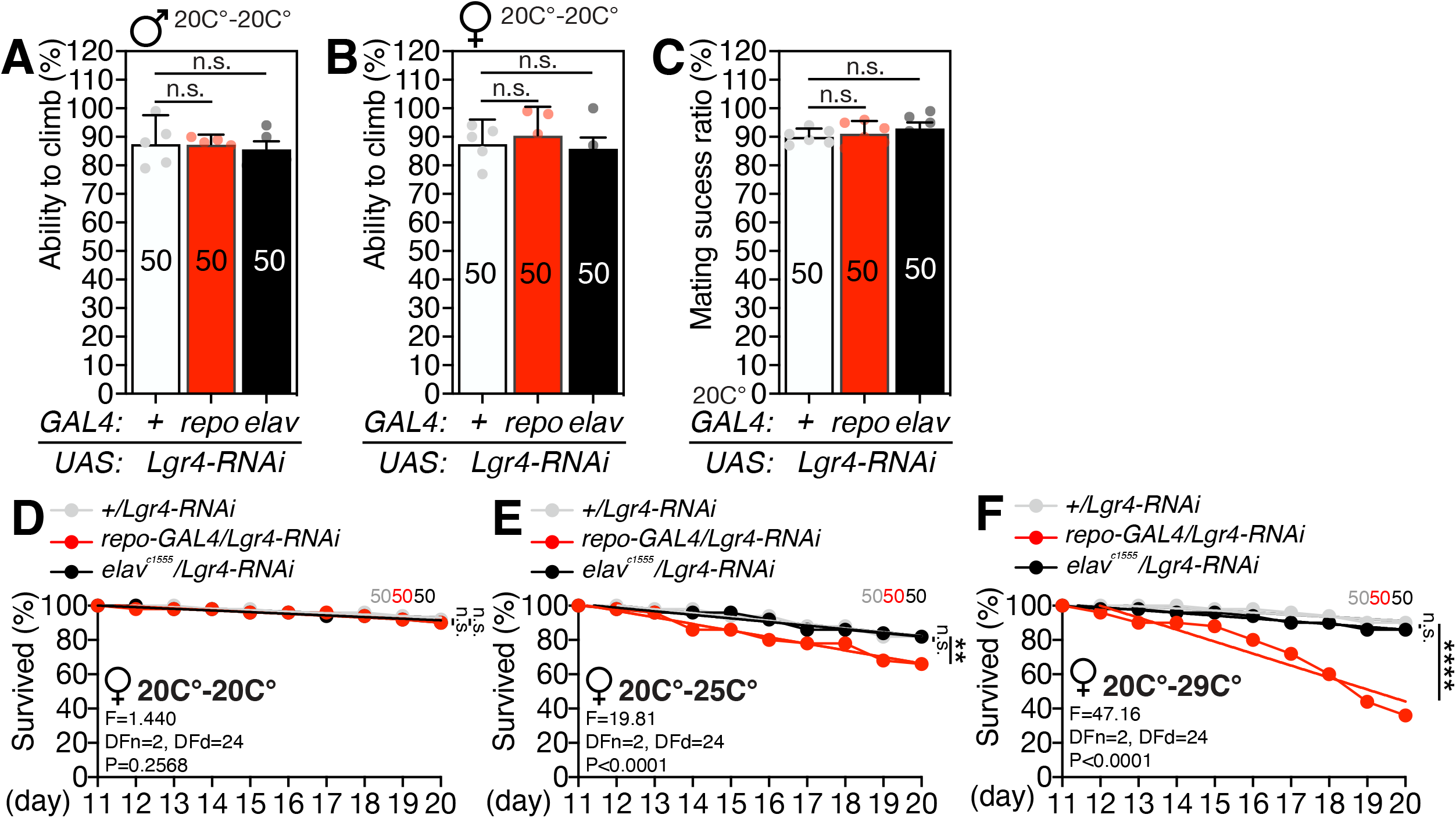
Control experiments for the impact of Lgr4 knockdown in adult behavior and physiology. (A-B) Climbing assay of (A) male or (B) female flies expressing *Lgr4-RNAi* in permissive temperature 20°C. (D-E) Lifespan assay of female flies expressing *Lgr4-RNAi* either in neuron or glia. Genotypes and rearing condition of all above data are labeled in the graph.

**Fig. S4.**
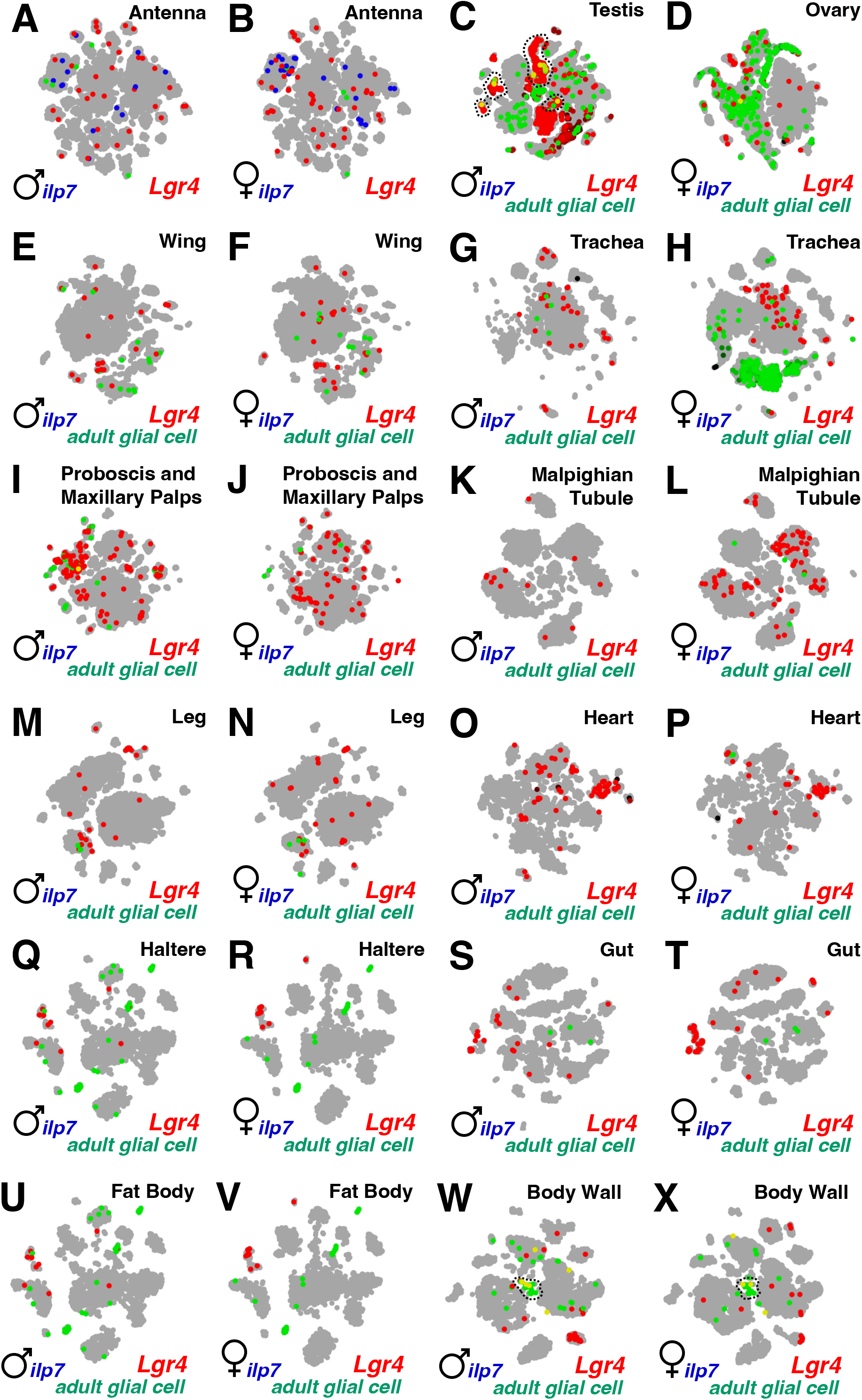
snRNAseq of adult fly in various tissues annotated by fly SCope. (A-X) SCope data for different annotated tissues. Tissues, sex, and gene names of all above data are color-coded using red, green, and blue words. When cells overlap, the color of the dots is either yellow, cyan, or magenta. Dotted lines indicate glial cell populations that expresses *Lgr4*.

**Fig. S5.**
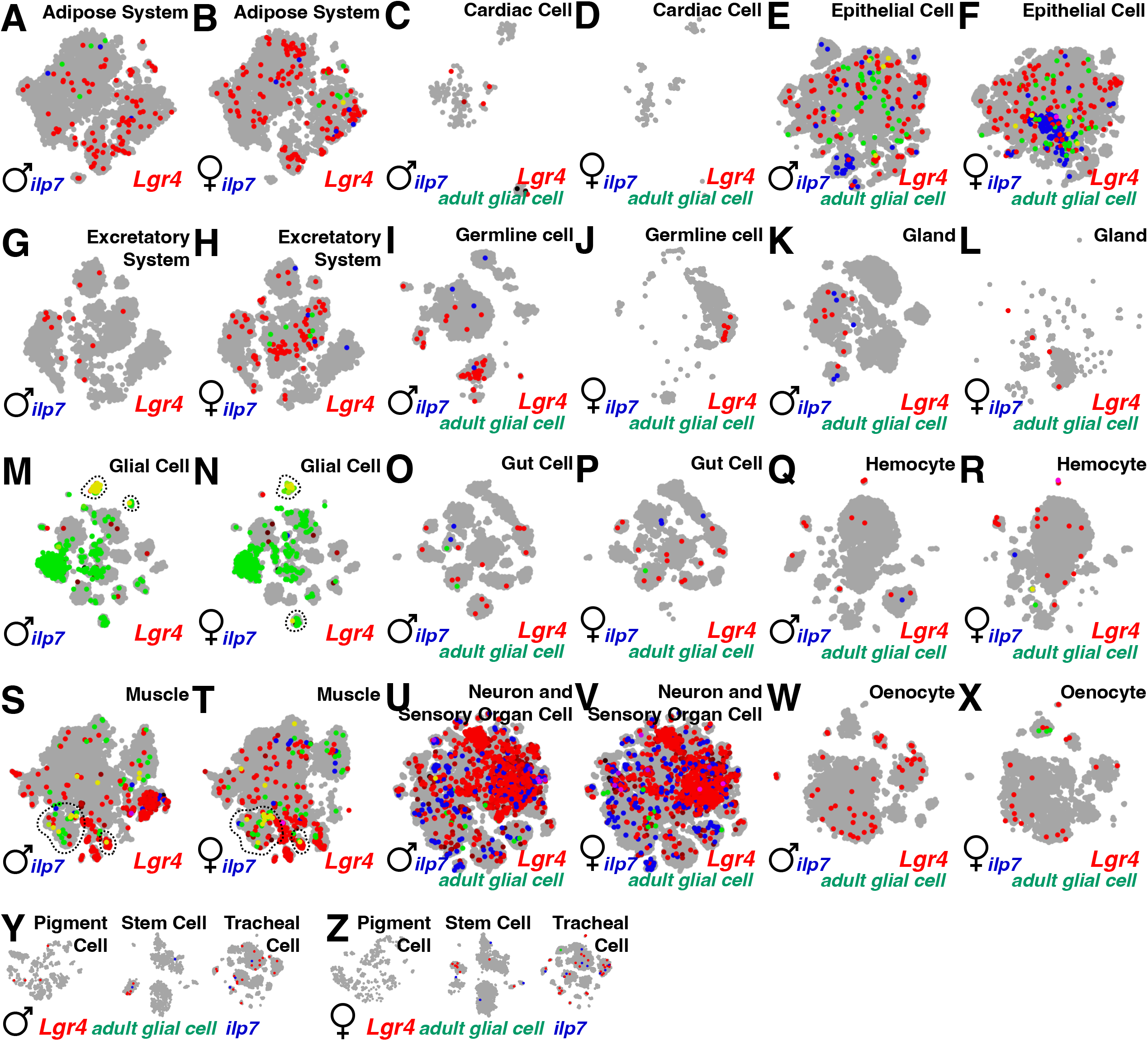
snRNAseq of adult fly in various cell populations annotated by fly SCope. (A-Z) SCope data for different annotated cell populations. Annotations, sex, and gene names of all above data are color-coded using red, green, and blue words. When cells overlap, the color of the dots is either yellow, cyan, or magenta. Dotted lines indicate glial cell populations that expresses *Lgr4*.

**Fig. S6.**
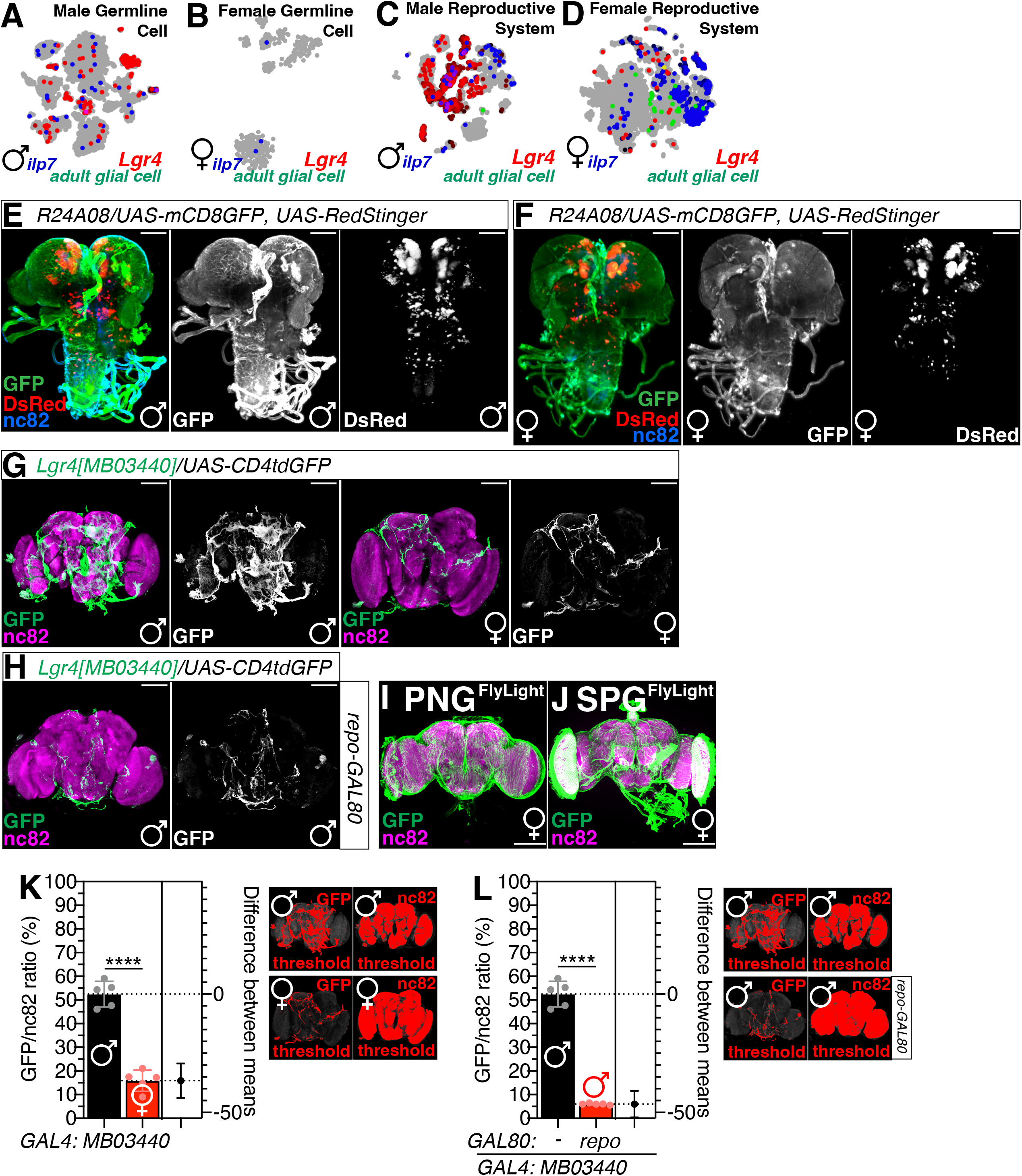
*Lgr4-GAL4s* label glia in larval and adult CNS. (A-D) SCope data for different annotated sex-specific tissues. (E-F) Third instar (B) male or (C) female larval CNS expressing *R24A08-GAL4* with *UAS-CD4tdGFP, UAS-RedStinger* were immunostained with anti-GFP (green), anti-DsRed (Red) and anti-nc82 (blue) antibodies. Gray panel represents GFP and RedStinger signals. Scale bars represent 100 μm. (G-H) Adult brains expressing (G) *Lgr4[MB03440]* or (H) with *repo-GAL80* with *UAS-CD4tdGFP* were immunostained with anti-GFP (green) and anti-nc82 (magenta) antibodies. Gray panel represents GFP signals for quantification to set threshold shown in the right panels of (K and L). (I-J) Adult brains expressing (I) *PNG-* or (J) *SPG-GAL4* drivers with *UAS-myrGFP*. Source image lsm files (~500MB) were downloaded from FlyLight platform constructed by Janelia Farm Research Center (JFRC) then reconstructed using ImageJ https://www.janelia.org/project-team/flylight. (K-L) Ratio of GFP/nc82 quantified from (G) and (H) to compare (K) male vs. female or (L) *GAL4* only vs. *GAL4* with *GAL80*. GFP signals represents *GAL4* expressing region and nc82 staining region were calculated by ImageJ threshold and measure function then quantified. See **EXPERIMENTAL PROCEDURES** and previous report [31,32] for detailed quantification methods.

## EXPERIMENTAL PROCEDURES

### Fly Rearing and Strains

*Drosophila melanogaster* were raised on cornmeal-yeast medium at similar densities to yield adults with similar body sizes. Flies were kept in 12 h light: 12 h dark cycles (LD) at 25°C (ZT 0 is the beginning of the light phase, ZT12 beginning of the dark phase) except for some experimental manipulation (experiments with the flies carrying *RNAi* reared in 20°C or 29°C for experimental purposes). Wild-type flies were Canton-S. To reduce the variation from genetic background, all flies were backcrossed for at least 3 generations to CS strain. All mutants and transgenic lines used here have been described previously.

We are very grateful to the colleagues who provided us with many of the lines used in this study. We obtained the following line from Dr. Marc Freeman (Vollum Institute, OHSU, USA): *NP2222-GAL4*; from Drs. Qingzhong Ren and Tzumin Lee (HHMI Janelia Research Campus, USA): *Eaat1^P88^-GAL4, Mz709-GAL4:* from Dr. Chun Han (Cornell University, USA) *A58-GAL4*.

The following lines were obtained from Bloomington Stock Center (#stock number): *UAS-dicer* (#24650, #24651), *TM3, y[+] Ser[1]/Sb[1]* (#1614), *Lgr4-RNAi* (#28655), *repo-GAL4* (#7415), *elav^c155^; UAS-Dcr-2* (#25750), *Lgr1-RNAi* (#51465), *Lgr2-RNAi* (#31958), *Lgr3-RNAi* (#55910), *alrm-GAL4* (#67032, 67031), *Eaat1-GAL4* (#8849), *GCM-GAL4* (#35541), *17A-GAL4* (#8474), *A8C-GAL4* (#35543), *Mz97-GAL4* (#9488), *ALG-GAL4* (#45914), *CG-GAL4* (#39944), *EGN-GAL4* (#39157), *EGT-GAL4* (#39908), *PNG-GAL4* (#40436), *SPG-GAL4* (#50472), *UAS-mCD8GFP* (#5130), *UAS-RedStinger* (#8547), *UAS-CD4tdGFP* (#35836), *Lgr4[MB01485]* (#23395), *Lgr4[MB23615]* (#23395), *R25C08-GAL4* (#23395), *R24A08-GAL4* (#23395); from Vienna Drosophila Stock Center (#stock number) [34]: *Lgr4-IR2* (#v102681), *Lgr4-IR3* (#v108915); from Kyoto Stock Center (#stock number): *NP2222-GAL4* (#112830).

### Lethality Assay

We evaluated the lethality of various *RNAi* strains expressed by tissue-specific GAL4s using the method we described elsewhere (Reference ALS2022). Briefly, to measure the effects of each cross for lethality, we used TM3 or TM6B balancer chromosomes. As shown in Fig. 1, S1, 2 and 3, virgin *RNAi/TM6B* female flies and male *GAL4/TM3* flies were crossed and the proportion of progeny with the *TM3* or *TM6B* balancer was counted and compared the progeny without *TM3* or *TM6B*. For control experiments shown in Fig. 1A-B, we used male *TM3, y[+] Ser[1]/Sb[1]* then measures the ratio of *Ser[1]* and *Sb[1]* marker phenotype.

### Climbing Assay

For climbing assay, we modified the conventional RING assay [35]. In brief, 40-50 aged flies were placed in an empty vial and were tapped to the bottom of the tube. We used 10 days old adults reared in different temperatures. After tapping of flies, we recorded 10 seconds of video clip. This experiment was done five times with 5-minute intervals. With recorded video files, we captured the position of flies 10 seconds after tapping the vial. This captured image file was then be loaded in ImageJ to perform particle analysis.

### Particle Analysis with Climbing Assay Data

For quantifying the location of flies inside vial, we used the “analyze particles” function of ImageJ [36]. The position of pixels was normalized by height of vial then only the particles above the midline (4 cm) of vial were counted.

### Lifespan Assay and Statistical Analysis

For lifespan analysis, we used conventional procedure [37]. Briefly, 50 flies were aged by sex before being raised in typical 12 h light: 12 h dark cycles at either 29°C, 25°C or 20°C for each experimental objective. The number of dead flies was recorded daily for forty days. Every three to four days, the surviving flies were transferred to fresh vials. To compare the survival curves of each genotype, we used the software Graph Pad Prism’s linear aggression analysis. When linear regression analysis reveals that the slopes of the survival curves across genotypes are significantly different, we consider the survival curves between genotypes to be distinct.

### Larval CNS Dissection and Immunostaining

As described before [38], wandering third instar larval CNS or adult brain was dissected then fixed in 4% formaldehyde for 30 min at room temperature, washed with 1% PBT three times (30 min each) and blocked in 5% normal donkey serum for 30 min. The brains were then incubated with primary antibodies in 1% PBT at 4°C overnight followed with fluorophore-conjugated secondary antibodies for 1 hour at room temperature. Brains were mounted with anti-fade mounting solution (Invitrogen, catalog #S2828) on slides for imaging. Primary antibodies: chicken anti-GFP (Aves Labs, 1:1000), rabbit anti-DsRed express (Clontech, 1:250) and mouse anti-Bruchpilot (nc82) (DSHB, 1:50). Fluorophore-conjugated secondary antibodies: Alexa Fluor 488-conjugated goat anti-chicken (Invitrogen, 1:100), RRX-conjugated donkey anti-rabbit (Jackson Lab, 1:100), and Dylight 649-conjugated donkey anti-mouse (Jackson Lab, 1:100).

### Quantitative Analysis of GFP Fluorescence

To quantify the GFP signals in larval and adult CNS, we measured fluorescence intensity using the measure tool of ImageJ (National Institutes of Health, http://rsb.info.nih.gov/ij) as described previously [31,32]. Fluorescence was quantified in a manually set region of interest (ROI). For quantifying fluorescence signals, we used the threshold and measure function of ImageJ.

### Statistical Analysis

Statistical analysis of climbing assay was similar with our previous studies [32,38,39]. 40-50 adults were used for climbing assay. Statistical comparisons were made between groups that were naively reared or sexually experienced within each experiment. As climbing data of adults showed normal distribution (Kolmogorov-Smirnov tests, p > 0.05), we used two-sided Student’s t tests. Each figure shows the mean ± standard error (s.e.m) (**** = p<0.0001, *** = p < 0.001, ** = p < 0.01, * = p < 0.05). All analysis was done in GraphPad (Prism). Individual tests and significance are detailed in figure legends. Besides traditional t-test for statistical analysis, we added estimation statistics for all climbing assays and two group comparing graphs. In short, ‘estimation statistics’ is a simple framework that—while avoiding the pitfalls of significance testing—uses familiar statistical concepts: means, mean differences, and error bars. More importantly, it focuses on the effect size of one’s experiment/intervention, as opposed to significance testing [40]. In comparison to typical NHST plots, estimation graphics have the following five significant advantages such as (1) avoid false dichotomy, (2) display all observed values (3) visualize estimate precision (4) show mean difference distribution. And most importantly (5) by focusing attention on an **effect size**, the difference diagram encourages quantitative reasoning about the system under study [41]. In 2019, the Society for Neuroscience journal eNeuro instituted a policy recommending the use of estimation graphics as the preferred method for data presentation [42]

### Single-nucleus RNA-sequencing Analyses - Data and Code Availability

snRNAseq dataset analyzed in this paper is published in [27] and available at the Nextflow pipelines (VSN, https://github.com/vib-singlecell-nf), the availability of raw and processed datasets for users to explore, and the development of a crowd-annotation platform with voting, comments, and references through SCope (https://flycellatlas.org/scope), linked to an online analysis platform in ASAP (https://asap.epfl.ch/fca).

